# A model-based high throughput method for fecundity estimation in fruit fly studies

**DOI:** 10.1101/382804

**Authors:** Enoch Ng’oma, Elizabeth G. King, Kevin M. Middleton

## Abstract

The ability to quantify fecundity is critically important to a wide range of experimental applications, particularly in widely-used model organisms such as *Drosophila melanogaster.* However, the standard method of manually counting eggs is time consuming and limits the feasibility of large-scale experiments. We develop a predictive model to automate the counting of eggs from images of eggs removed from the media surface and washed onto dark filter paper. A cross-validation approach demonstrates our method performs well, with a correlation between predicted and manually counted values of 0.88. We show how this method can be applied to a large data set where egg densities vary widely.

## Introduction

Reproductive output, with its close tie to Darwinian fitness, is potentially the most important of an individual’s phenotypes and thus a critical phenotype to be able measure in many different experiments. While counting the number of eggs or offspring produced by an individual is simple in concept, in practice it is often quite challenging. In many insect systems, including the widely used model organism *Drosophila melanogaster (D. melanogaster)*, females produce a large number of eggs over their lifetime, and the most common method of quantifying lifetime fecundity is by manually counting eggs soon after they are laid. This manual counting is often achieved by visualizing the media surface under a dissecting scope and counting each egg on the surface, often using a grid to keep track of already counted areas and using tools such as a hand-held tally counter to keep an accurate count. Alternatively, the media itself may be digitally imaged or eggs may be washed onto a dark surface and then imaged, with the number of eggs in the image counted by looking at a zoomed in image and tallying each egg. These features limit the ability to quantify lifetime fecundity for large numbers of experimental replicates. Thus, the development of methods that allow the automation of egg counting while maintaining accuracy have the potential to expand the set of biological questions that can be investigated.

As image analysis techniques in general have gotten more sophisticated, there is a growing interest in applying these techniques to different types of biological data^1^. There are several challenges associated with applying image analysis methods to fecundity data in *D. melanogaster*. Female flies lay eggs on the food media, which for most standard recipes^2^ is close in color to the eggs and does not provide a high contrast background. This challenge has been addressed in the past by providing egg laying media that is higher contrast (e.g., transparent media^3^ or the addition of charcoal to the media^4^), or by washing the eggs off of the surface of the food and filtering them through black filter paper^5–7^. Irregularities on the surface of the food, the presence of dust or other particulate matter, and the clumping of many eggs together are all issues that have the potential to affect any automated image analysis. Existing methods for automating egg counting in *Drosophila* have developed methods to identify individual eggs on the food surface and then quantify the total number present^3,4^. While these methods perform well for some applications, their performance declines when eggs are at high densities with a lot of clumping, or when there are many other similarly sized non-egg objects in the image. In addition, because both rely on images of the food surface, for recipes that produce lightly colored food, which may be necessary in studies with a specific diet manipulation, it becomes more difficult for these methods to accurately quantify eggs.

We sought to develop a method for robustly predicting egg counts applicable to a wide range of applications, with a minimal impact of the challenges described above. Our method first produces images with a high contrast between eggs and the background and then takes advantage of the simple relationship between the area of light-colored pixels and the number of eggs present to develop a predictive model that can be applied to a large set of images. This method has the flexibility to be applied to a wide range of egg densities and can be applied to many experimental contexts.

## Materials and methods

### Fecundity samples

The set of samples used to develop and validate our egg counting method were part of a quantitative genetic study employing a half-sibling, split family design, with families split over three different dietary conditions using flies derived from the *Drosophila* Synthetic Population Resource^8^. All vials began with 15-24 female flies and 6 male flies with fecundity estimated once per week over a 24-hour oviposition period for the entire lifetime of the flies. Females were provided with fresh media each Monday and following a 24-hour egg laying period, these egg vials were collected the following Tuesday and stored at −20 ° C until processing. We continued this process each week until all females within a vial had died. To visualize eggs, we modified a protocol developed by Rose and colleagues^5–7^. We added 2.5 mL of 50% bleach to each vial (frozen or thawed) and gently swirled at 20 orbits per minute for about 2 min 50 s on an ORBITRON Rotator 1 (Boekel Scientific, USA) to separate eggs from the media. Eggs were flushed from each vial onto a black filter disc fit into a custom-built vacuum apparatus with a brief spray of embryo solution (12% Triton X-100, SIGMA-ALDRICH, USA) followed by a spray of water to rinse. As a quality check, flask filtrate was re-filtered to a new disc periodically, and processed vials inspected under a dissecting microscope. The filter discs were prepared from black landscape fabric (DEWITT Weed Barrier Pro, www.dewittcompany.com) to a radius of 3.65 cm.

Discs bearing eggs were photographed using a Canon EOS Rebel T5i (Canon Inc., Japan) camera vertically attached to a steel clamp using the following settings: exposure time 1/25 seconds, aperture F5.6, and ISO 100. Egg-bearing discs were mounted on a black stage placed within a cubical photo studio 35cm x 35cm x 35cm constructed with a PVC frame and enclosed by a white fabric. The entire setup was placed in a dark room lit by two facing white light lamps, 23W, 1200 lumens (model # LBP23PAR2/5K, UTILITECH, China) clamped to the stage outside the studio box to allow for soft uniform lighting across the imaging platform. Image files were immediately inspected for quality with ImageJ^9^.

### Estimation of egg counts

We developed a predictive model to estimate the counts of eggs from the images of the filter discs. Figure 1 gives a graphical overview of our methodology for acquiring and processing images (Fig. 1a) and the development of the model (Fig. 1b) described below. To create the model and assess its performance, eggs on 345 discs were manually counted from the images of the discs using the *Multi-point* tool in ImageJ^9^ to generate a training dataset. This tool allows the user to mark each egg in an image by clicking on its position and automatically keeps a tally of the total number marked in an image. To ensure that all possible imaging conditions were available for model training, discs selected for manual egg counting were selected across the whole lifespan of flies and across all three diet treatments. Some background reflectivity of the discs and small debris present meant that some non-zero white area was present in almost all images, so our training set includes an over-representation of images with relatively few eggs, to prevent upward bias in the fecundity predictions.

**Figure 1:**
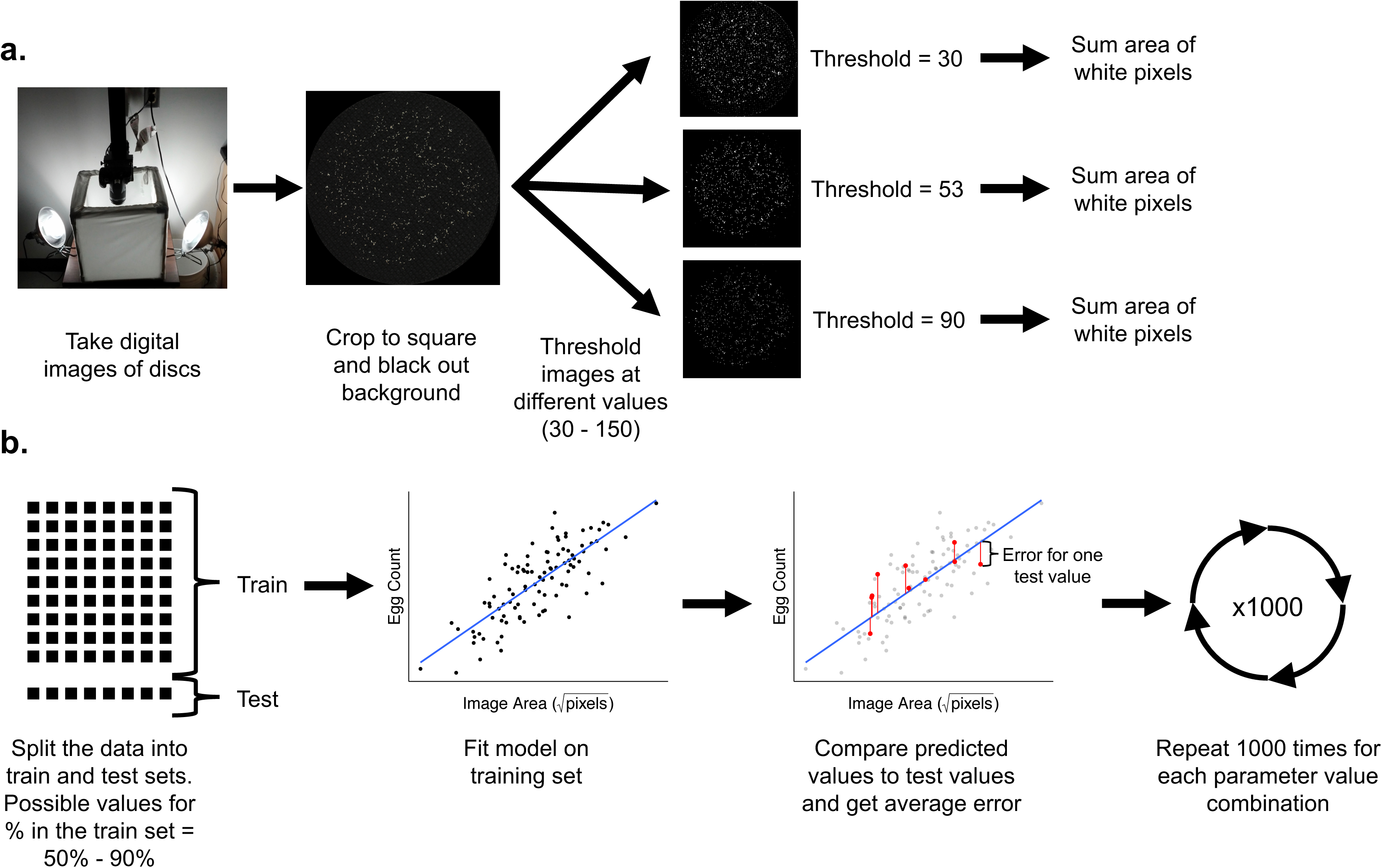
Graphical overview of the fecundity prediction method. **a.** Image acquisition and processing steps. **b.** Model fitting and optimization steps. See Methods for full details.

Each image was manually cropped to be square at the edges of the disc, and the background outside the disc was set to black to eliminate any reflectivity or debris occurring outside of the disc using GNU Image Manipulation Program^10^. Subsequently, the image was binarized by thresholding 8-bit grayscale values using the OpenCV package^11^ in Python^12^ (v. 3.6; https://github.com/EGKingLab/fecundity_estimation). From this thresholded image, the area of white pixels was summed (https://github.com/EGKingLab/fecundity_estimation). Because images were not all the same size due to slight variations in camera zoom, we included size of the image as a covariate in the linear model predicting egg count from white pixel area and image dimension:

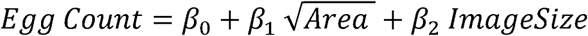

where *Area* is the sum of the white pixels and *Image Size* is the linear dimension of one edge of the image in pixels. We use 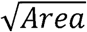 because pixel area is predicted to scale as a square of egg count.

The optimal grayscale threshold value was determined by cross-validation^13^ using R^14^ (v. 3.5 https://github.com/EGKingLab/fecundity_estimation). We tested threshold grayscale values between 30 and 150 (for 8-bit images with 256 grayscale levels), across a range of training/testing data splits (0.5 to 0.9) and a range of proportions of data (0.5 to 1.0). In cross-validation, a dataset is split into training and test sets. The training set is a portion of the data used to create the predictive model. Thus, in this case, the training set is used to fit the linear model described above. The test set is used to assess the ability of the model to predict new data (i.e., data not included to produce the predictive model). In this case the model fit with the training set would produce a predicted egg count from white area and image size, and that value is compared to the manual count to quantify the error in prediction for new data. We also varied the proportion of the total set of manually-counted images we used to develop the model in order to test whether we were using a large enough set of manually counted images to appropriately train the model. Each model was fit 1,000 times at each combination of threshold, train/test split, and data proportion, with the root mean squared difference between predicted count and manual count (RMSd) retained for each iteration (https://github.com/EGKingLab/fecundity_estimation). The optimal set of parameters was determined from the mean RMSd of these 1,000 iterations (Fig. 2). That threshold was a grayscale value of 53 for the training set, using 90% of the data each iteration for training, and only required using 50% of the total data in any one iteration. The resulting predictive model performed well for our training set (Fig. 3a). The trained model was then used to predict egg counts for all 3,768 images that were part of this dataset (Fig. 3b). Images with negative predicted egg counts were set to 0. All images were also visually inspected, and those determined to be of poor quality, with excessive background reflectivity, were counted manually and were not included in the training or prediction set.

**Figure 2:**
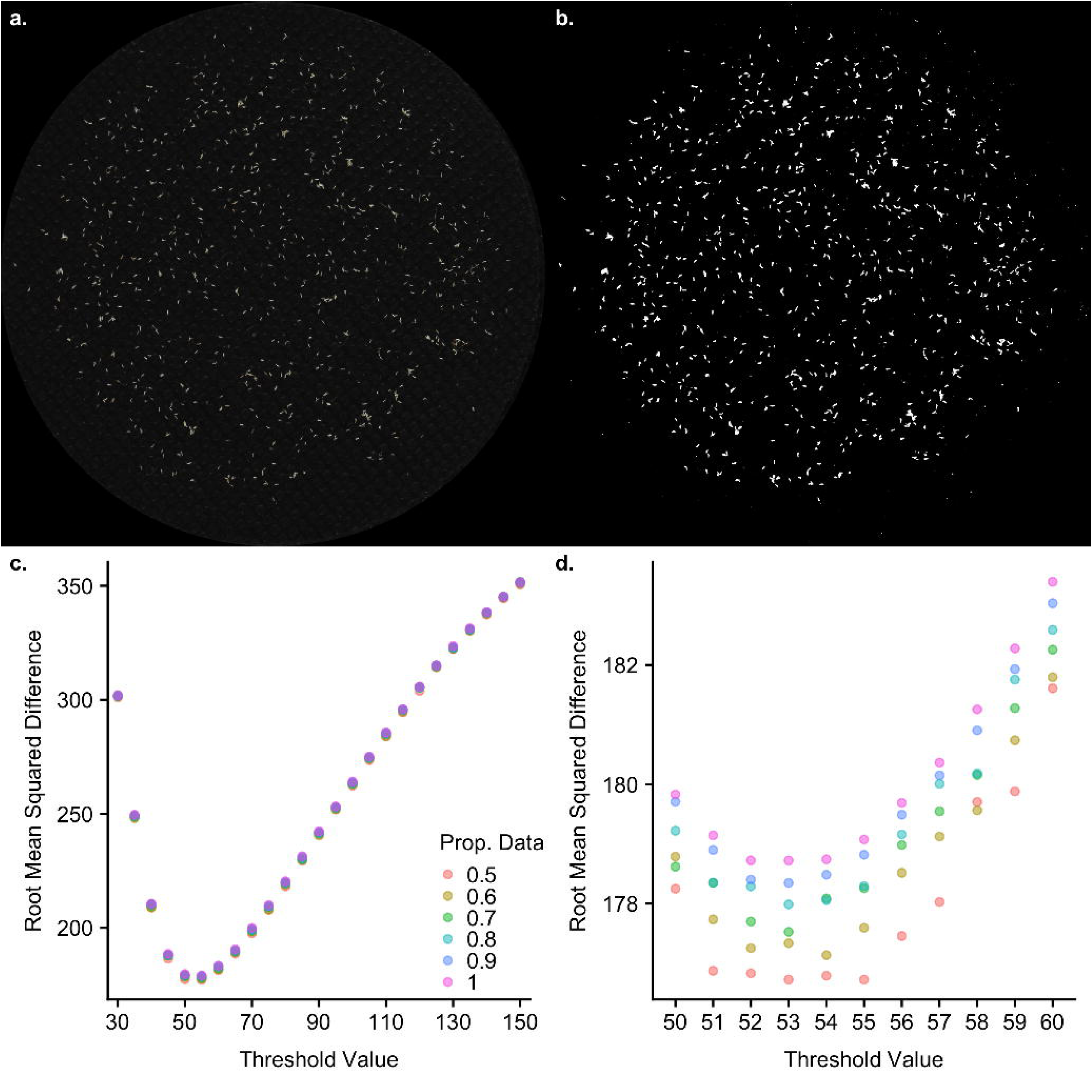
Performance of the model predicting egg counts from images. **a.** A representative raw image of eggs separated from fly media and filtered onto black filter fabric. **b.** The same image thresholded at the optimal threshold level identified by the model. Due to the time required for optimization of threshold value, we carried this process out in two steps. **c**. Coarse optimization was carried out for every 5 grayscale values between 30 and 150, which revealed a single minimum in that range, falling between 50 and 60. **d**. Fine optimization was then carried for integer values between 50 and 60, resulting in a single optimal value of 53. Colors represent the proportion of data used for each model fit. Even at the optimal value, the difference in minimum (at 50% of data) and maximum (at 100% of data) in root mean squared difference was ∼2 eggs.

**Figure 3:**
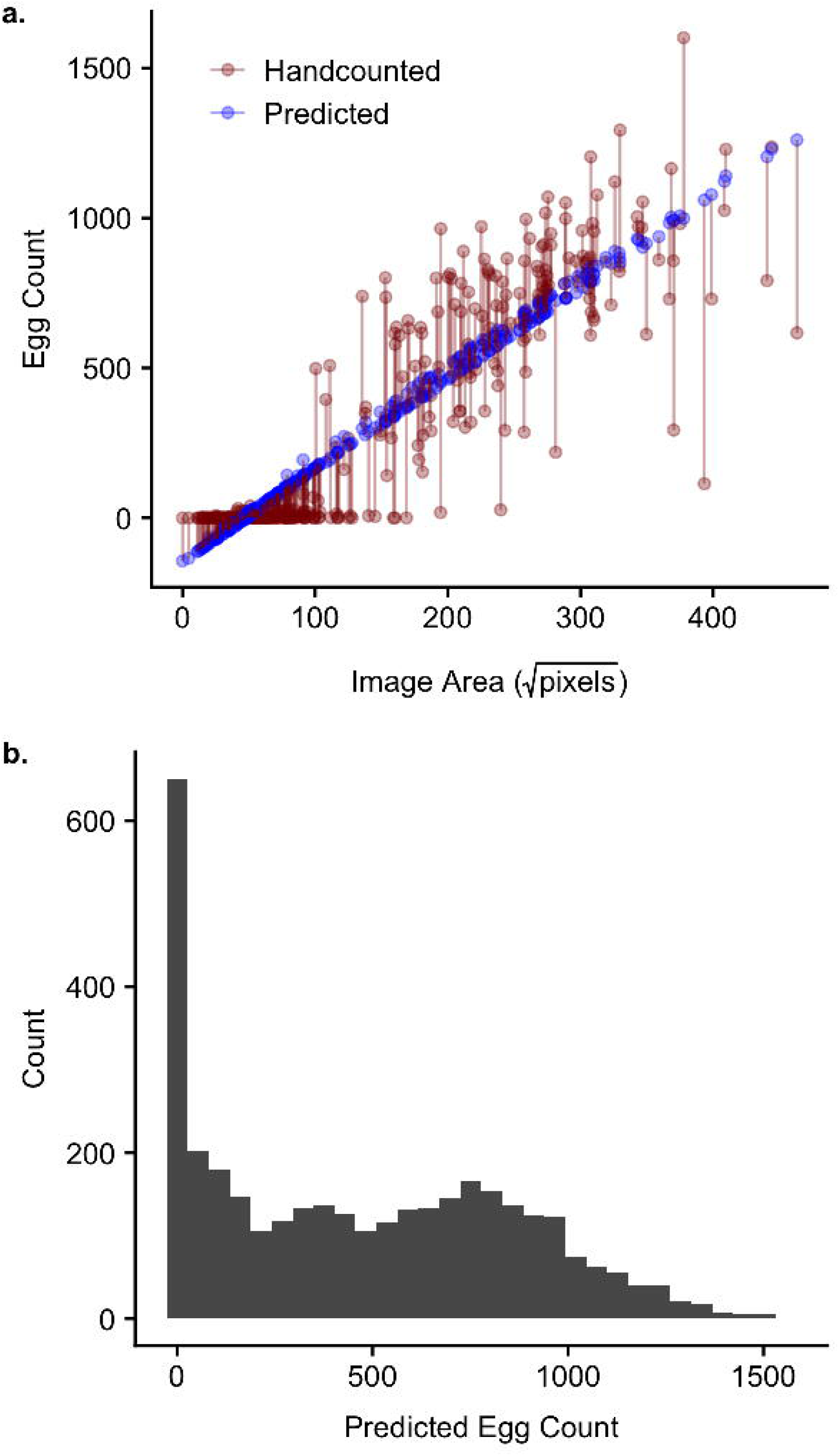
The performance of the predictive model. **a.** The square root of image area is on the x-axis and the egg count is on the y-axis. True, manually counted values are shown in red and the predicted values from the model are shown in blue. Pairs of values (manually counted and predicted) for individual images are connected by a red line. **b**. Distribution of predicted values for all images in the data set. Inflated zero counts results from assigning a value of 0 to any predicted values <0.

### Data and software availability

Our raw data, including our fecundity estimates, and all egg image files, have been uploaded to Zenodo and may be retrieved at http://doi.org/10.5281/zenodo.1285237. All scripts to reproduce all of our analyses are available at GitHub: https://github.com/EGKingLab/fecundity_estimation. Code is available to directly download egg image files from the Zenodo and reproduce our analyses.

## Results

We developed a high throughput method for obtaining egg counts by creating a predictive model to obtain estimated egg counts from images of eggs that were washed off the food surface and filtered onto a dark fabric disc by exploiting the simple linear relationship between the number of eggs present and the amount of white area on a thresholded image of a disc containing *D. melanogaster* eggs (Fig. 2a). We used a set of manually counted images to optimize the model and show that it performs well (Fig 3a).

We tested several parameters to produce an optimal prediction model. First, by using different proportions of our total dataset of manually counted images, we showed our manually counted set was sufficiently large to train the predictive model and achieve a high predictive ability. Using only 50% of the manually counted data, we achieved similar error rates in the resulting model, demonstrating our dataset was sufficiently large to saturate the model (Fig 2c, d). We also tested a range of thresholds to identify the value that would remove the most background noise without obscuring actual eggs (Fig 2c, d). To assess the robusticity of our model across different parameters, we employed cross-validation to ensure we tested the ability of the model to predict egg counts for new data (data not used to fit the model). For each iteration, 90% of the data was used to train the model and we assessed the ability of this model to predict the values of the remaining 10% of the data. Our best performing model was able to predict new data well, with a mean absolute error of ∼177 eggs and a high correlation (*r* = 0.89) between the predicted egg count and the manual egg counts. We note that with a 24-hour egg laying period for a vial with several females, this error rate, with predicted counts deviating from manual counts by an average of 177 eggs, represents a relatively small number of eggs. After identifying the optimal threshold value, we fit a model using our full dataset and found similar error rates when comparing the predicted values from this model to the manually counted images (*r* = 0.88, mean absolute error = 128 eggs; Fig. 3a). We were then able to use this model to predict egg counts for our entire dataset of 3,768 images (Fig. 3b).

## Discussion

The method of washing eggs from the surface of the food onto black filter paper has been used previously in studies quantifying fecundity in *D. melanogaster*^5–7^. While this is more time consuming than imaging the surface of the food directly, this approach has several advantages. First, removing eggs from the surface of the food typically results in less clumped eggs and removes surface irregularities such as bubbles in the food. Second, it allows for eggs to be imaged on a high contrast background no matter what food recipe is used. This feature is particularly important when diet is a manipulated variable in an experiment and/or it is undesirable to add charcoal or some other darkening agent to the food. Finally, any imaging (including on the food surface) provides a permanent record of the phenotype while manually counting and discarding vials does not allow for validation of the phenotyping or later error checking. In previous work using this method, eggs washed onto black filter paper were still manually counted. Here, we have shown how this process can be automated, providing a significant time saving, e.g., avoiding manual counts of ∼84,000 eggs in this study.

Other automated egg counting methods rely on identifying individual objects in an image as eggs, including those of Waithe et al.^3^ and Nouhaud et al.^4^ It is not surprising then that both methods perform more poorly when egg densities are high and multiple eggs clump together. The method we have developed is able to handle a wide range of densities because the egg count is estimated directly from the area of white pixels, removing the need to identify eggs as individual objects. The method described here is susceptible to debris on the filter paper surface (e.g., food particles, etc.), because any lightly colored object will be included in the total thresholded area. We included a large number of low egg count discs to help calibrate the average amount of background debris on our discs. Nevertheless, it is important when applying this method to make sure minimal amounts of debris are present. This may be achieved either by careful egg-washing and/or pre-cleaning discs before imaging (and recording eggs that lie on the debris being removed).

Overall, we present an approach to automating egg counting that can be optimized and tailored to a wide variety of situations and applications. We use a cross-validation, train/test approach to identify the model parameters that allow us to optimally predict new egg count values. The cross-validation approach shows a high correspondence between predicted and manually counted egg counts for new data (data not used to form the predictive model), showing our high performance is not inflated due to overfitting to our manually counted set. This approach could potentially be applied to any images with enough contrast to threshold the background from the objects of interest and may be especially applicable to situations where it is challenging to separate individual objects.

## Acknowledgements

Larry Cabral provided with valuable ideas through personal communication to establish our fecundity protocol. Elizabeth Lopresti, Michael Reed, Osvaldo Enriquez, Anna Perinchery and Kyla Winford helped with fly husbandry, time-intensive experimental set-ups, collection and entry of data. This work was supported by NIH grant R01 GM117135 to Elizabeth G. King, The University of Missouri, and a University of Missouri Research Board Grant.

## References

1. Nketia TA, Sailem H, Rohde G, Machiraju R, Rittscher J. Analysis of live cell images: Methods, tools and opportunities. Methods 2017; 115:65–79.

2. Bloomington Drosophila Stock Center. Fly Food Recipes [Internet]. 2017 [cited 2018 Aug 1]; Available from: https://bdsc.indiana.edu/information/recipes/index.html

3. Waithe D, Rennert P, Brostow G, Piper MDW. QuantiFly: Robust Trainable Software for Automated Drosophila Egg Counting. PLoS ONE 2015; 10:e0127659.

4. Nouhaud P, Mallard F, Poupardin R, Barghi N, Schlötterer C. High-throughput fecundity measurements in Drosophila. Sci Rep [Internet] 2018 [cited 2018 Jul 25]; 8. Available from: https://www.ncbi.nlm.nih.gov/pmc/articles/PMC5849729/

5. Rose Lab Data. Fecundity: Animated Introduction [Internet]. 2012; Available from: https://www.youtube.com/watch?v=xrEBNB8bUwo&sns=em

6. Rose Lab Data. Fecundity: Live Demonstration [Internet]. 2012; Available from: https://www.youtube.com/watch?v=uqFZ4ZTC6ZY&sns=em

7. Burke MK, Barter TT, Cabral LG, Kezos JN, Phillips MA, Rutledge GA, Phung KH, Chen RH, Nguyen HD, Mueller LD, et al. Rapid divergence and convergence of life-history in experimentally evolved Drosophila melanogaster. Evolution 2016; 70:2085–98.

8. Ng’oma E, Fidelis W, Middleton K, King EG. The evolutionary potential of diet-dependent effects on lifespan and fecundity in a multi-parental population of Drosophila melanogaster. bioRxiv 2018; 343947.

9. Rueden CT, Schindelin J, Hiner MC, DeZonia BE, Walter AE, Arena ET, Eliceiri KW. ImageJ2: ImageJ for the next generation of scientific image data. BMC Bioinformatics 2017; 18:529.

10. The GIMP Team. GNU Image Manipulation Program [Internet]. 2015. Available from: http://www.gimp.org

11. Bradski, G. The OpenCV Library. Dr Dobb’s Journal of Software Tools 2000;

12. Python Software Foundation. Python Language Reference [Internet]. Available from: http://www.python.org/

13. James G, Witten D, Hastie T, Tibshirani R, editors. An introduction to statistical learning: with applications in R. New York: Springer; 2013.

14. R Core Team R: A language and environment for statistical computing. R Foundation for Statistical Computing [Internet] 2018; Vienna, Austria. Available from: http://www.R-project.org/

